# A role for the circadian transcription factor NPAS2 in the progressive loss of non-rapid eye movement sleep and increased arousal during fentanyl withdrawal in male mice

**DOI:** 10.1101/2022.04.27.489767

**Authors:** Mackenzie C. Gamble, Byron Chuan, Teresa Gallego-Martin, Micah A. Shelton, Stephanie Puig, Christopher P. O’Donnell, Ryan W. Logan

## Abstract

**Rationale:** Synthetic opioids like fentanyl are contributing to the rise in rates of opioid use disorder and drug overdose deaths. Sleep dysfunction and circadian rhythm disruption may worsen during opioid withdrawal and persist during abstinence. Severe and persistent sleep and circadian alterations are putative factors in opioid craving and relapse. However, very little is known about the impact of fentanyl on sleep architecture and sleep-wake cycles, particularly opioid withdrawal. Further, circadian rhythms regulate sleep-wake cycles, and the circadian transcription factor, neuronal PAS domain 2 (NPAS2) is involved in the modulation of sleep architecture and drug reward. Here, we investigate the role of NPAS2 in fentanyl-induced sleep alterations.

**Objectives:** To determine the effect of fentanyl administration and withdrawal on sleep architecture, and the role of NPAS2 as a factor in fentanyl-induced sleep changes.

**Methods:** Electroencephalography (EEG) and electromyography (EMG) was used to measure non-rapid eye movement sleep (NREMS) and rapid eye movement sleep (REMS) at baseline and following acute and chronic fentanyl administration in wild-type and NPAS2-deficient male mice.

**Results:** Acute and chronic administration of fentanyl led to increased wake and arousal in both wild-type and NPAS2-deficient mice, an effect that was more pronounced in NPAS2-deficient mice. Chronic fentanyl administration led to decreased NREMS, which persisted during withdrawal, progressively decreasing from day 1 to 4 of withdrawal. The impact of fentanyl on NREMS and arousal was more pronounced in NPAS2-deficient mice.

**Conclusions:** Chronic fentanyl disrupts NREMS, leading to a progressive loss of NREMS during subsequent days of withdrawal. Loss of NPAS2 exacerbates the impact of fentanyl on sleep and wake, revealing a potential role for the circadian transcription factor in opioid-induced sleep changes.

## Introduction

Fentanyl is a highly potent synthetic opioid used to alleviate pain in certain clinical contexts including during and after surgery and in the treatment of cancer and chronic pain (Comer and Cahill 2019). Fentanyl along with other synthetic opioids have surpassed heroin and oxycodone as the main drivers of overdose deaths and increased rates of opioid use disorder (OUD) in regions with already high base rates of opioid use (Gladden 2016; O’Donnell 2017). Thus, understanding the unique consequences of fentanyl use on health is imperative to developing strategies for treatments to curb the impact. An understudied health-related impact of chronic opioid use has been at the intersection of sleep and circadian rhythms.

Accumulating evidence suggests opioids lead to sleep and circadian disruptions, possibly contributing to increased craving and propensity to relapse (for review see Eacret et al. 2020; Logan et al. 2014). Sleep abnormalities are evident in OUD (Wang and Teichtahl 2007; Fathi et al. 2020) and likely contribute to relapse (Lydon-Staley et al. 2017). The standard treatment, methadone, for OUD and opioid dependence also leads to notable sleep disturbances (Tripathi et al. 2020). For example, morphine, heroin, and methadone acutely suppress rapid eye movement sleep (REMS) and promote wakefulness in humans (Kay et al. 1969; Lewis et al. 1970; Kay 1975). However, as tolerance to repeated use of opioids develops, the impact on REMS lessens with levels approaching individual baselines (Lewis et al. 1970; Kay 1975). Non-rapid eye movement sleep (NREMS) is also disrupted by opioids (Kay et al. 1969; Orr and Stahl 1978; Shaw et al. 2005). Notably, withdrawal from opioids leads persistent disruptions in both NREMS and REMS, possibly lasting for months in patients (Oswald 1969; Lewis et al. 1970; Kay 1975). Similarly, in animal models, both NREMS and REMS are altered during and following opioid administration and withdrawal (Khazan and Colasanti 1972; De Andrés and Caballero 1989). In addition, disrupted sleep impacts opioid tolerance and withdrawal. For example, sleep deprivation delays the development of morphine tolerance in rats (Ahmadi-Soleimani et al. 2021) and reduces opioid-induced analgesia in humans (Smith et al. 2020). Further understanding of the complex relationships between sleep, circadian rhythms, and opioids is essential for advancing effective treatments for OUD and opioid dependence.

To begin to fill this gap, we investigated the impact of chronic fentanyl administration and withdrawal in mice on sleep architecture. We investigated the effects of acute and chronic fentanyl, along with withdrawal from fentanyl, on NREMS and REMS in male mice. Additionally, we explored the potential role for the circadian transcription factor, neuronal PAS domain 2 (NPAS2), in the modulation of opioid-induced sleep changes. NPAS2 is preferentially expressed in the mammalian forebrain, an essential component of the molecular clock, and previously been shown to regulate drug reward (Ozburn et al. 2015) and sleep architecture (Dudley et al. 2003). NPAS2 is uniquely positioned at the intersection of sleep, circadian rhythms, and opioids. Therefore, we used both wild-type (Wt) and NPAS2-deficient (Garcia et al. 2000) male mice to investigate fentanyl-induced changes on sleep architecture.

## Methods

### Animals and Housing

Experiments were conducted in Wt and NPAS2-deficient male mice (Garcia et al. 2000). Littermates were used as controls. Mice were initially group-housed then transferred to individual cages at least five days prior to surgical procedures (n=7 per group per genotype). Mice were housed under standard 12:12-hour light-dark schedule (lights on at 08:00h and off at 20:00h) with *ad libitum* food (Prolab RMH 3000 5P76, LabDiet) and water. During sleep recordings, mice were placed into a pyramidal cage to allow for the recording cable to be tethered to the mouse and connected to the recording equipment via a commutator. Mice were then placed into secondary housing container to insulate noise and other stimuli during sleep recordings. Experiments were conducted in accordance with the Guide for the Care and Use of Laboratory Animals of the National Institutes of Health and approved by the Institutional Animal Care and Use Committee at the University of Pittsburgh School of Medicine.

### Sleep Polysomnography Surgery

Mice were implanted with electroencephalogram (EEG) and electromyogram (EMG) electrodes, as previously described (Tagaito et al. 2001). Briefly, an incision along the midline of the head was used to expose the skull and muscles immediately posterior to the skull. Underlying fascia was then gently cleared from the skull and holes (1mm diameter) were drilled through the skull (frontal and parietal). EEG electrodes (E363/1, Plastics One Inc.) were secured via screws into the holes (first electrode at 2-3mm caudal to Bregma and 1-2mm right of the midline; second electrode at 2-3mm rostral to bregma and 2-3 right of the midline; and third electrode 0-1mm rostral to bregma and 2-3mm left of midline. EMG electrodes (E363/76, Plastics One Inc.) were then secured onto the surface of the muscle immediately posterior to the dorsal portion of the skull. EEG and EMG electrodes were inserted into a pedestal (MS363, Plastics One Inc.) and cemented to the skull with dental acrylic (1403 and 1420, Lang Dental). Skin caudal to the pedestal was sutured. Mice recovered for five days following surgery.

### Sleep Polysomnography Recording

Mice were tethered to a preamplifier system via a custom cable (363-SL/6 80CM 6TCS, Plastics One Inc.) for sleep recordings. Our sleep-wake detection system has previously been described and validated in mice (Benington et al. 1994; O’Donnell et al. 2019). Sleep-wake states were determined using the frequency distribution of EEG and the amplitude of EMG. Temporal data was considered in terms of “epochs” of 10s duration. Software thresholds using the polysomnographic parameters EEG frequency and EMG amplitude were set to assess sleep-wake state (Tagaito et al. 2001). For EEG, thresholds for β2 (20–30Hz) and δ1 (2–4Hz) from the EEG frequency distribution were used to parse NREMS (lower β2/δ1 ratio) from wakefulness or REMS (above β2/δ1 threshold). For EMG, above the highest threshold represented wakefulness (irrespective of the β2/δ1 ratio), while below the lowest threshold represented either NREMS or REMS depending on whether the β2/δ1 ratio was below (NREMS) or above (REMS) their threshold. The amplitude from EEG was also used to determine REMS, such that the amplitude needed to be below a threshold. Amplitude was included because in mice this measure invariably undergoes a uniform reduction during the transition from NREMS to REMS. Thresholds were optimized for each mouse using baseline EEG and EMG recordings (1-2 hours).

### Sleep-Wake State Scoring

Continuous EEG and EMG data was recorded as waveforms using WinDaq Data Acquisition Software (v.2.94, DATAQ Instruments, Inc.) and converted into an analyzable format (Stanford Sleep Structure Scoring System) via a custom program (Benington et al. 1994), subsequently validated in mice (Veasey et al. 2004). Manual sleep-wake state validation was based on previously described parameters (O’Donnell et al. 2019). Time spent in NREMS, REMS, and awake states were calculated as the percent of total spent in that state during a complete 24- hour period. Bouts of NREMS and REMS were based on the following criteria: 1) NREMS bout began with more than three consecutive epochs of NREM and ended with either more than three consecutive epochs of wake or more than two consecutive epochs of REMS; 2) REMS bout began with more than two consecutive epochs of REMS and ended with more than three consecutive epochs of NREMS or wake; and 3) wakefulness bout began with more than three consecutive epochs of wake and ended with more than three consecutive epochs of NREMS and/or REMS. Propensity to remain awake was defined as the average duration from the beginning of more than three consecutive epochs of wake to the following of more than three consecutive epochs of NREMS and/or REMS over 24 hours. Arousal index was defined as the number of arousals per hour of sleep over 24 hours, with an arousal being more than a single epoch of wake following more than three epochs of NREMS and/or REMS. Independent investigators involved in the study separately scored sleep state for each mouse. Analyses presented were completed using the mean values between the scorers.

### Fentanyl Administration and Sleep Recordings

Male Wt and NPAS2-deficient mice underwent either acute or repeated administration of fentanyl (320µg of fentanyl in sterile saline per kg body weight administered intraperitoneal, i.p.). For mice receiving acute fentanyl, a single injection of fentanyl was administered on day 12 after receiving 11 previous days of saline (**Fig. 1**). For mice receiving repeated fentanyl, fentanyl was injected every 12 hours (08:00h and 20:00h) from days 3 to 10 (**Fig. 1**). Sleep was recorded for all mice on day 0, prior to any saline or fentanyl administration. Sleep was also recorded on days 13 and 14 to determine the acute effects of fentanyl on sleep and during acute withdrawal (first 24 hours following last fentanyl administration). Sleep was recorded on days 10 to 14 to determine the effects of repeated fentanyl on sleep and during prolonged withdrawal (initial four days following last fentanyl administration).

**Fig. 1.**
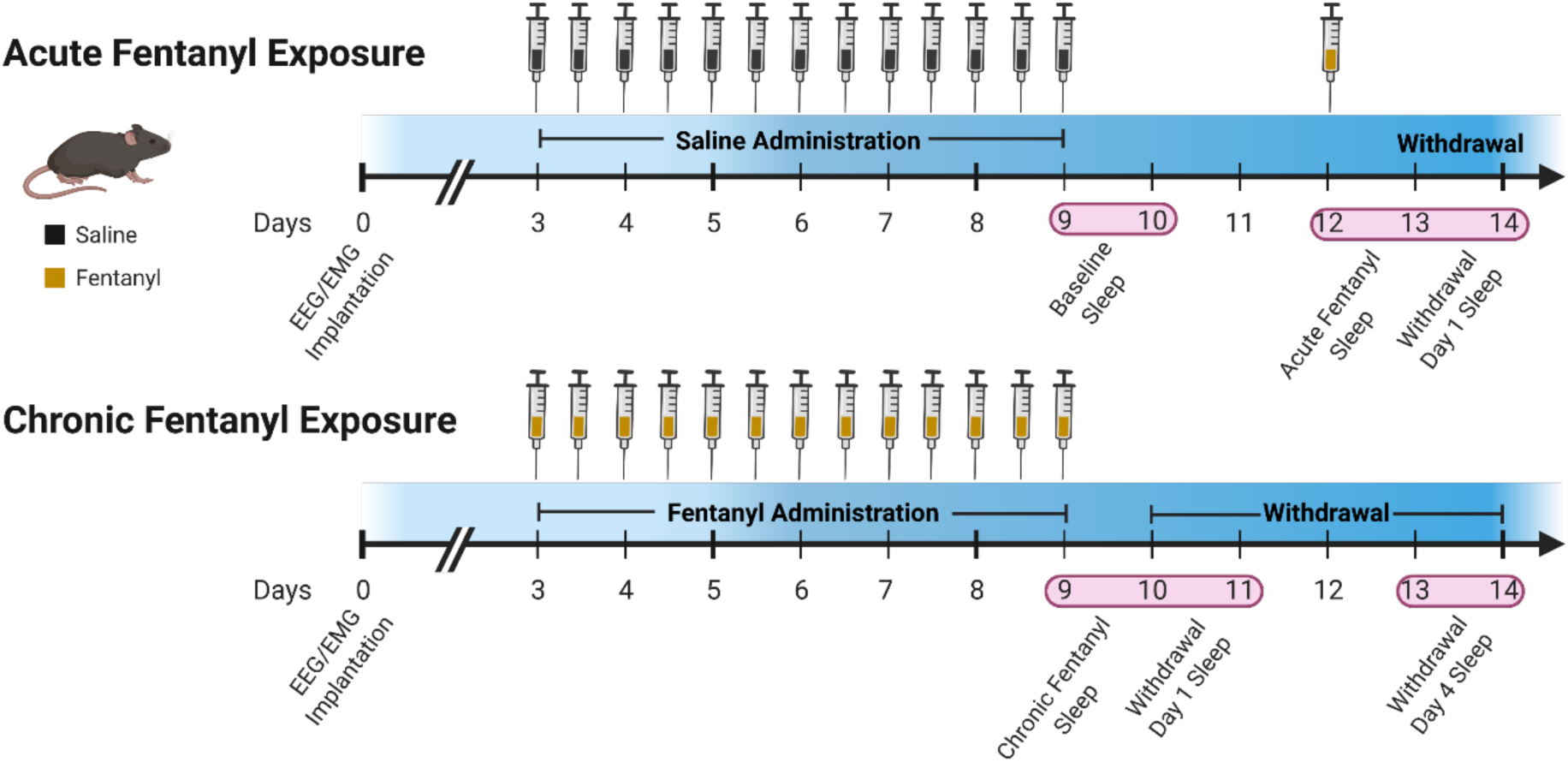
Schematic of fentanyl administration in mice. Wild-type and NPAS2-/- mice underwent electroencephalogram (EEG) and electromyography (EMG) implantation to record sleep-wake states, followed by chronic administration with saline (black) or fentanyl (orange) via i.p. injection twice a day for 7 days. Sleep-wake state recordings assessed sleep architecture at baseline, and during acute and chronic fentanyl administration, followed by withdrawal. Created using BioRender.

### Statistical Analyses

Two-way mixed ANOVA (between within design) or Mixed Effects Model when unbalanced was used to compare changes in sleep states between and NPAS2-deficient mice among acute and chronic fentanyl administration groups, and during fentanyl withdrawal across time. Dunnett’s (experimental groups compared to control group), Sidak (repeated measure with two levels), and Tukey’s post-hoc (repeated measure with three levels) analyses were used to test multiple comparisons. Geisser-Greenhouse correction was used for all applicable analyses. Outliers were determined using the three-sigma rule. This included 1 outlier in sleep bouts and sleep bout length measures for baseline, acute fentanyl administration, chronic fentanyl administration, and acute withdrawal in NPAS2-deficient mice. Results are presented as means ± standard error of the mean (SEM) and significance was set at α=0.05.

## Results

### Effects of saline administration on sleep and arousal in wild-type and NPAS2-deficient mice

Sleep was measured using EEG/EMG in mice following saline and fentanyl administration (**Fig. 1**). Baseline NREMS and REMS was measured immediately following repeated administration of saline. Overall, the amount of time in each sleep stage were similar between Wt (mean ± standard error of mean over 24 hrs for NREMS: 51.8 ± 1.36% and REMS: 3.3 ± 0.23%, and NPAS2-deficient (NREMS 49.83 ± 1.80% 24hr and REMS 2.917 ± 0.35%, **Fig. 2A,B**) mice. In addition, the time to transition from sleep to wake (arousal index) was similar between Wt (19.4 ± 1.18 minutes) and NPAS2-deficient male mice (26.75 ± 3.40 minutes, **Fig. 2C**). Time to fall asleep (sleep propensity) was also similar (Wt: 2.77 ± 0.30 minutes to fall asleep after ≥3 wake epochs over 24 hrs compared to NPAS2-deficient: 2.38 ± 0.32 minutes, **Fig. 2D**). The number of sleep bouts and length of bouts were largely comparable between Wt (98.71 ± 8.68 bouts over 24 hrs with an average length of 8.54 ± 0.66 minutes, **Fig. 2E**) and NPAS2-deficient (92.17 ± 6.97 bouts over 24 hrs with an average length of 8.77 ± 0.52 minutes (**Fig. 2F**). Overall, baseline sleep measures of NREMS, REMS, bouts, and bout durations were similar between male Wt and NPAS2-deficient mice.

**Fig. 2.**
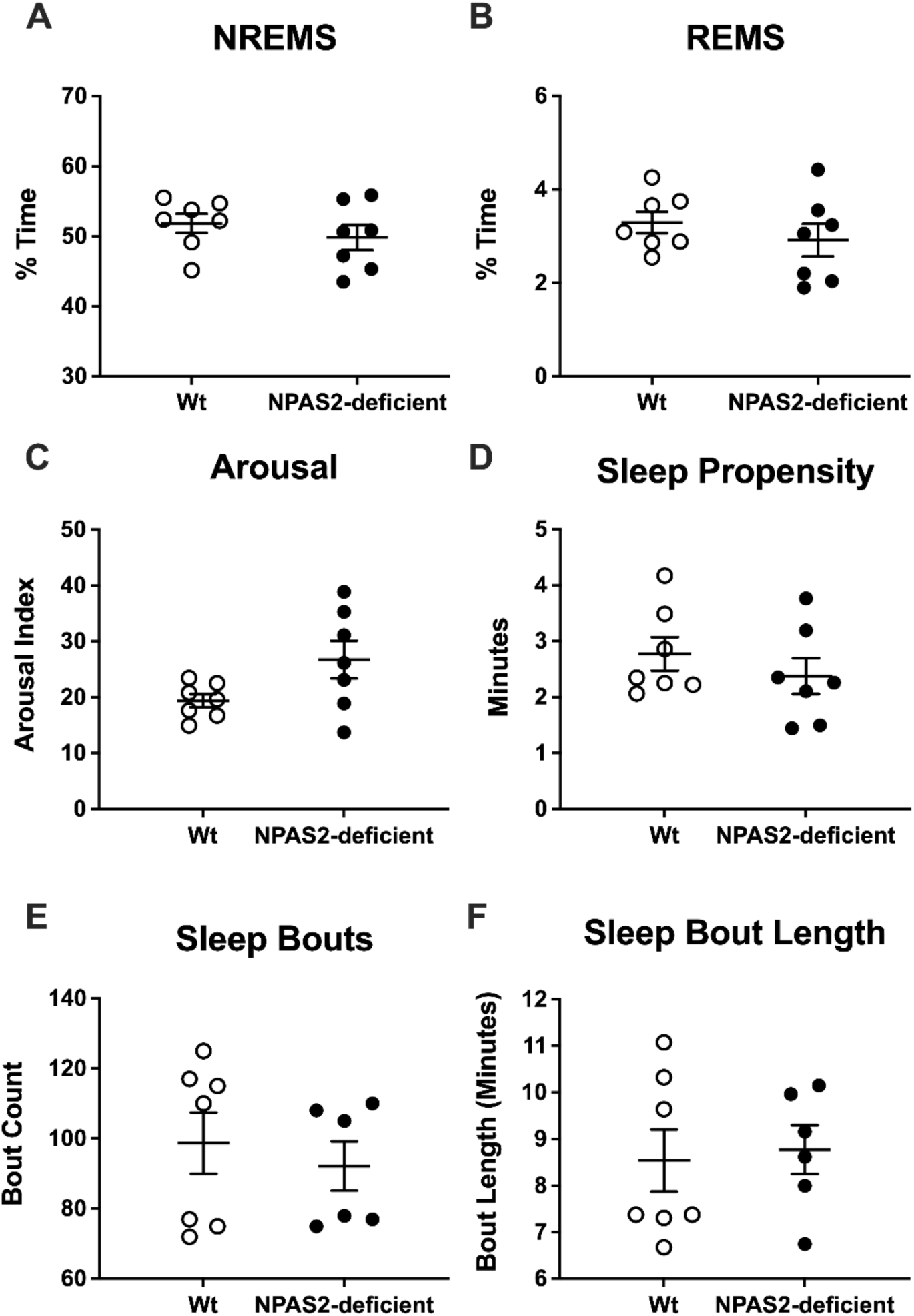
Similar sleep-wake states between wild-type and NPAS2-deficient male mice. Sleep-wake analysis of wild-type (Wt) and NPAS2-deficient mice at baseline after 7 days of saline treatment. Sleep-wake state was recorded for a 24 hr duration following the last saline injection on Day 9. **A.** Time spent in non-rapid eye movement sleep (NREMS) over 24h. **B.** Time spent in rapid eye movement sleep (REMS) over 24 hrs. **C.** Average number of minutes to transitions from sleep to wake state per hour of sleep over 24 hrs. **D.** Time to fall asleep after ≥3 awake epochs over 24 hrs. **E.** Sleep bouts over 24 hrs. **F.** Sleep bout length over 24 hrs.

### Effects of acute fentanyl on sleep and arousal in wild-type and NPAS2-deficient mice

To assess the impact of acute fentanyl on sleep, mice were administered an injection of fentanyl (320 ug/kg) following baseline recordings of sleep then sleep was measured for another 24 hrs (**Fig. 1**). Acute administration of fentanyl led to significant disruptions in sleep irrespective of genotype. Two-way ANOVA showed a main effect of acute fentanyl on NREMS (*F*_(1,12)_ = 11.8; *p* < 0.01) and no significant effect of genotype (p=0.06) or an interaction, along with no effect on REMS. NREMS was significantly decreased following acute fentanyl (from 50.84 ± 1.12% at baseline to 46.61 ± 1.42% after acute fentanyl collapsed across genotypes, **Fig. 3A,B**). Opioid-induced alterations of NREMS was attributed to a pronounced reduction displayed in NPAS2- deficient mice (from 49.83 ± 1.80% at baseline to 43.56 ± 1.76% after acute fentanyl, **Fig. 3A**). The duration of sleep to wake transitions (arousal index) were reduced by fentanyl (main effect of treatment, *F*_(1,12)_ = 9.2; *p* < 0.05, **Fig. 3C**), whereas sleep propensity significantly increased from 2.57 ± 0.22 minutes at baseline to 3.45 ± 0.26 minutes following acute fentanyl (main effect of treatment, *F*_(1,12)_ = 7.99; *p* < 0.05, **Fig. 3D**). The main effect of treatment for sleep propensity was mainly driven by an increase in NPAS2-deficient mice (from 2.38 ± 0.32 at baseline to 3.58±0.35 minutes over 24 hrs following acute fentanyl, **Fig 3D**). Total number of sleep bouts was significantly decreased from 95.69 ± 5.53 bouts across 24 hrs to 75.92 ± 4.82 bouts across genotypes (main effect of treatment, *F*_(1,12)_ = 6.92; *p* < 0.05, **Fig. 3E**), although no effects were observed for duration of sleep bouts (**Fig. 3F**). Thus, in the first 24 hrs following administration, acute fentanyl led to altered sleep, primarily in measures of arousal, sleep-wake transitions, and NREMS in both Wt and NPAS2-deficient male mice.

**Fig. 3.**
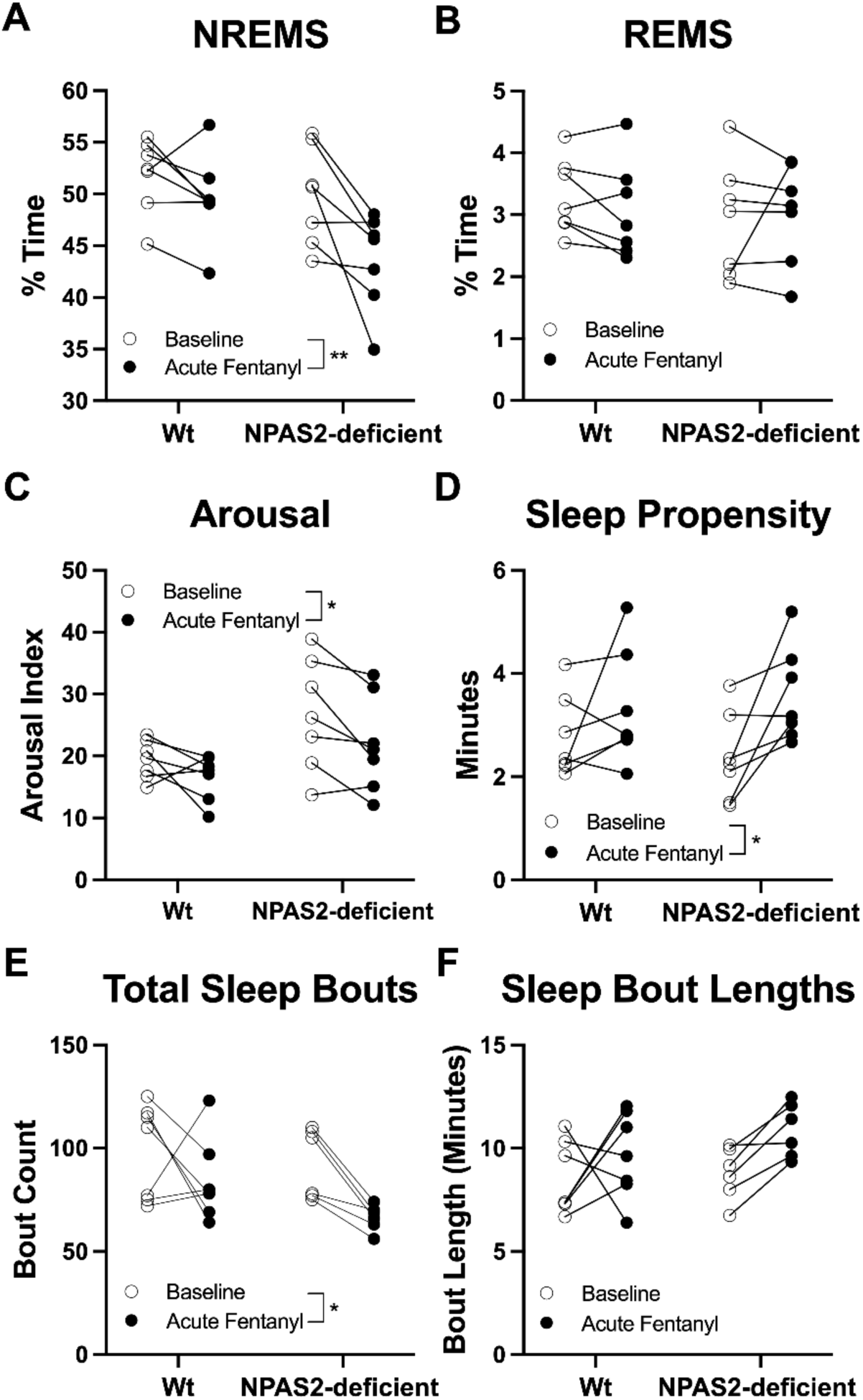
Acute fentanyl administration leads to altered sleep and arousal in male mice. Sleep- wake analysis of wild-type (Wt) and NPAS2-deficient mice at baseline compared to acute fentanyl administration. **A.** Time spent in non-rapid eye movement sleep (NREMS) over 24 hrs (main effect of treatment, \*\**p* < 0.01). **B.** Time spent in rapid eye movement sleep (REMS) over 24 hrs. **C.** Average number of transitions from sleep to wake state per hour of sleep over 24 hrs (main effect of treatment, \**p* < 0.05). **D.** Time to fall asleep after ≥3 awake epochs over 24 hrs (main effect of treatment, \**p* < 0.05). **E.** Total number of sleep bouts over 24 hrs (main effect of treatment, \**p* < 0.05). **F.** Average length of sleep bouts over 24 hrs.

### Effects of acute fentanyl withdrawal on sleep and arousal in wild-type and NPAS2-deficient mice

Sleep was also measured during the acute withdrawal period 24-48 hrs following acute administration of fentanyl. Acute fentanyl administration led to an overall reduction in NREMS that remained during the acute withdrawal period mainly in NPAS2-deficient mice (saline: 52.60 ± 1.55% across 24 hrs; acute fentanyl: 43.56 ± 1.76%; and acute withdrawal: 46.13 ± 1.94%, **Fig. 4A**). Two-way ANOVA showed a main effect of fentanyl condition (*F*_(1.78, 21.42)_ = 13.78; *p* < 0.001) and no significant effect of genotype (p=0.053) or an interaction on NREMS, along with no effects on REMS (**Fig. 4B**) an arousal index (**Fig. 4C**). Sleep propensity was altered by condition (main effect of treatment, *F*_(1.89, 22.79)_ = 5; *p* < 0.05, **Fig. 4D**). Overall, duration to fall asleep increased from baseline (2.67 ± 0.21 minutes) to 3.45 ± 0.26 minutes following acute fentanyl, and finally to 3.56 ± 0.23 minutes during withdrawal. For sleep bouts, a mixed-effects model (outlier removal, repeated measures) revealed main effects for genotype (*F*_(1,34)_ = 5.8; *p* < 0.05) and treatment (*F*_(1.65, 28.06)_ = 6.64; *p* < 0.01), reflected by an overall reduction in total sleep bouts by acute fentanyl administration that remained during the acute withdrawal period (**Fig. 4E**). An overall main effect of treatment was also found for sleep bout length (*F*_(1.84, 20.26)_ = 6.13; *p* < 0.01), whereby duration of sleep bouts were increased from baseline through acute withdrawal (8.86 ± 0.47 minutes per bout of sleep across 24 hrs during baseline, which increased to 10.21 ± 0.50 minutes after acute fentanyl administration, and further increased to 11.21 ± 0.53 minutes during acute withdrawal, **Fig. 4F**). Collectively, these findings indicate acute fentanyl led to alterations in sleep and arousal that were maintained during the acute withdrawal period similar across genotypes of mice, potentially reflecting the emergence of persistent changes in sleep.

**Fig. 4.**
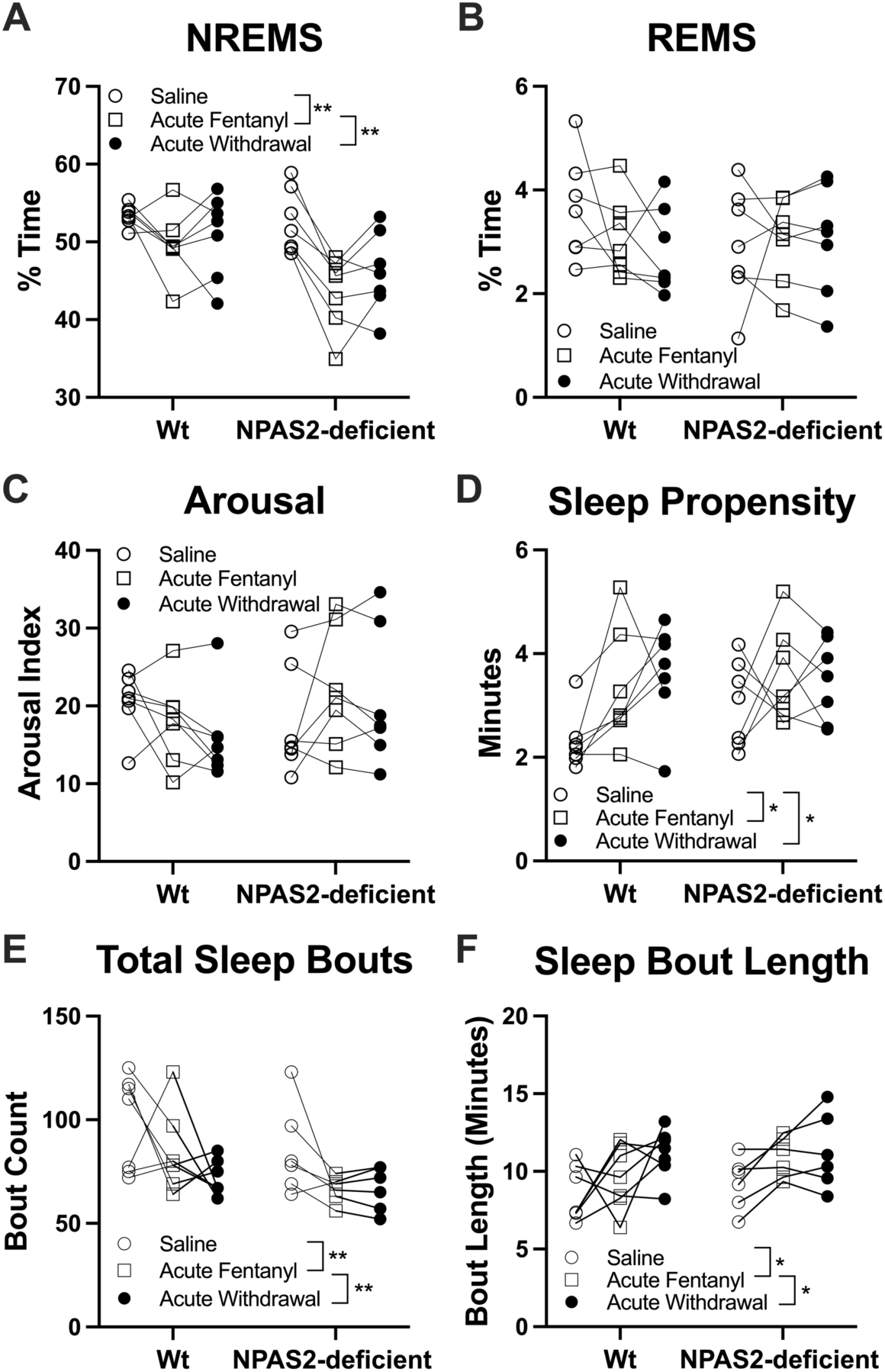
Acute fentanyl withdrawal leads to altered sleep and wake states that persist during acute withdrawal. Wild-type (Wt) and NPAS2-deficient mice after saline injection, 24 hrs following acute fentanyl administration, and 24-48 hrs after fentanyl (acute withdrawal). **A.** Time spent in non-rapid eye movement sleep (NREMS; saline vs. acute fentanyl, \*\**p* < 0.01; acute fentanyl vs. acute withdrawal, \*\**p* < 0.01). **B.** Time spent in rapid-eye movement sleep (REMS). **C.** Average number of arousals (transition from sleep to wake state) per hour of sleep (Tukey’s posthoc, saline vs. acute withdrawal in Wt mice, \**p* < 0.01). **D.** Time to fall asleep after ≥3 awake epochs (saline vs. acute fentanyl, \**p* < 0.05; saline vs. acute withdrawal, \**p* < 0.05). **E.** Total number of sleep bouts over the 24 hr recordings (saline vs. acute fentanyl, \*\**p* < 0.01; acute fentanyl vs. acute withdrawal, \*\**p* < 0.01). **F.** Average length of sleep bouts (saline vs. acute fentanyl, \**p* < 0.05; saline vs. acute withdrawal, \**p* < 0.05).

### Effects of chronic fentanyl administration and withdrawal on sleep and arousal in wild-type and NPAS2-deficient mice

In a separate cohort of mice, sleep was measured following chronic, repeated administration of fentanyl (7 days). In the initial 24 hrs following chronic administration of fentanyl, both NREMS and REMS were not significantly altered within and across genotypes of mice (**Fig. 5A,B**). A significant interaction was found for arousal index (genotype x treatment, *F*_(1,10)_ = 5.27; *p* < 0.05), with Sidak’s post hoc analyses indicating a significant reduction in NPAS2-deficient mice (baseline: 26.75 ± 3.40 minutes to transition from sleep to wake per hour of sleep over 24hrs on average to 17.75 ± 2.61 minutes, **Fig. 5C**). No significant effects were observed for sleep propensity (**Fig. 5D**). However, both the number (*F*_(1, 21)_ = 13.09; *p* < 0.01) and duration of sleep bouts (*F*_(1, 21)_ = 8.09; *p* < 0.01) were significantly altered by chronic fentanyl administration (**Fig. 5E,F**). The effects on sleep bouts were similar between acute (**Fig. 4E,F**) and chronic (**Fig. 5E,F**) among Wt and NPAS2-deficient mice, suggesting fentanyl led to an overall reduction in sleep bouts, while increasing bout duration in first 24-48 hrs following administration.

**Fig. 5.**
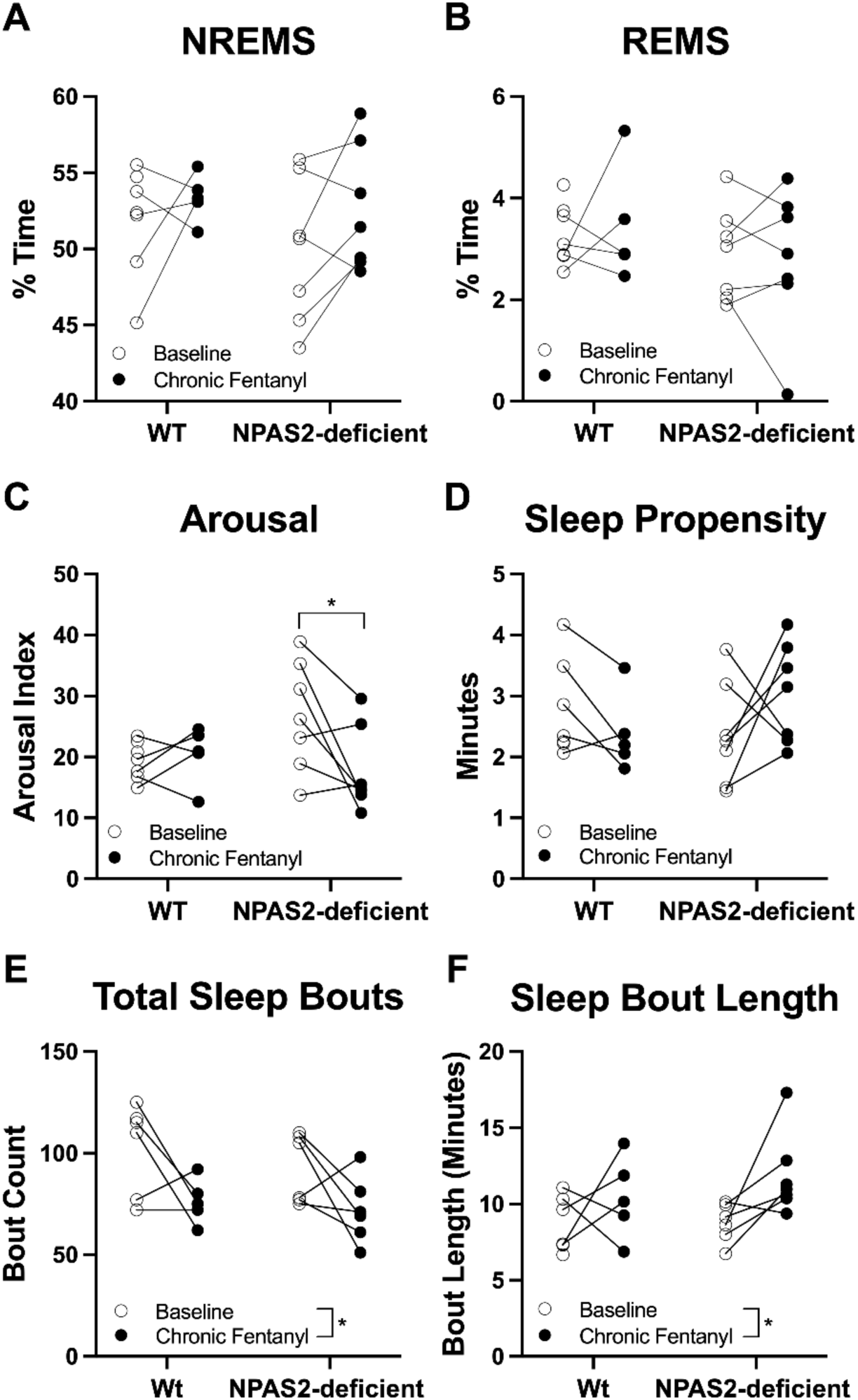
Chronic fentanyl administration leads to altered sleep and wake states. Wild-type (Wt) and NPAS2-deficient mice at baseline and after seven days of chronic fentanyl administration. **A.** Time spent in non-rapid eye movement sleep (NREMS) during 24 hrs of baseline compared to 24 hrs immediately following the final fentanyl administration (Day 9). **B.** Time spent in rapid-eye movement sleep (REMS). **C.** Average number of arousals (transition from sleep to wake state) per hour of sleep (baseline vs. chronic fentanyl in NPAS2-deficient mice, Sidak post hoc test, \**p* < 0.05). **D.** Time to fall asleep after ≥3 awake epochs. **E.** Total number of sleep bouts over 24 hrs (main effect of treatment, \**p* < 0.05). **F.** Average length of sleep bouts (main effect of treatment, \**p* < 0.05). Comparisons are between unpaired mice as the baseline and chronic fentanyl mice are from separate groups.

To determine whether opioid-induced alterations persisted during withdrawal, sleep was measured on days one and four of withdrawal. Both Wt and NPAS2-deficient mice exhibited a significant decrease in time spent in NREMS (main effect of treatment, F_(1.614,16.14)_ = 38.95; *p* < 0.0001) (**Fig. 6A**). In Wt mice, NREMS was 53.36 ± 0.69% after chronic fentanyl administration, which decreased to 50.96 ± 1.05% during the first day of withdrawal, then further decreased to 44.89 ± 0.75% after day four of withdrawal (∼9% reduction in NREMS). Similarly, NREMS progressively decreased in NPAS2-deficient mice during withdrawal from fentanyl (chronic fentanyl: 52.60 ± 1.55%; day 1: 48.45 ± 0.77%; and day 4: 44.29 ± 1.30%), evident by an ∼8% reduction in NREMS (**Fig. 6A**). However, inconsistent changes were observed in REMS between fentanyl administration and withdrawal from fentanyl regardless of NPAS2-deficiency (**Fig. 6B**). Further, while arousal was also unchanged, sleep propensity was significantly altered by fentanyl withdrawal (main effect of treatment, F_(1.127,11.27)_ = 18.90; *p* < 0.001), accompanied by an overall increase in both Wt and NPAS2-deficient mice (**Fig. 6D**). There were no significant changes in number of sleep bouts and duration of sleep bouts in Wt and NPAS2-deficient mice after either Day 1 or Day 4 of withdrawal from chronic fentanyl administration (**Fig. 6E,F**).

**Fig. 6.**
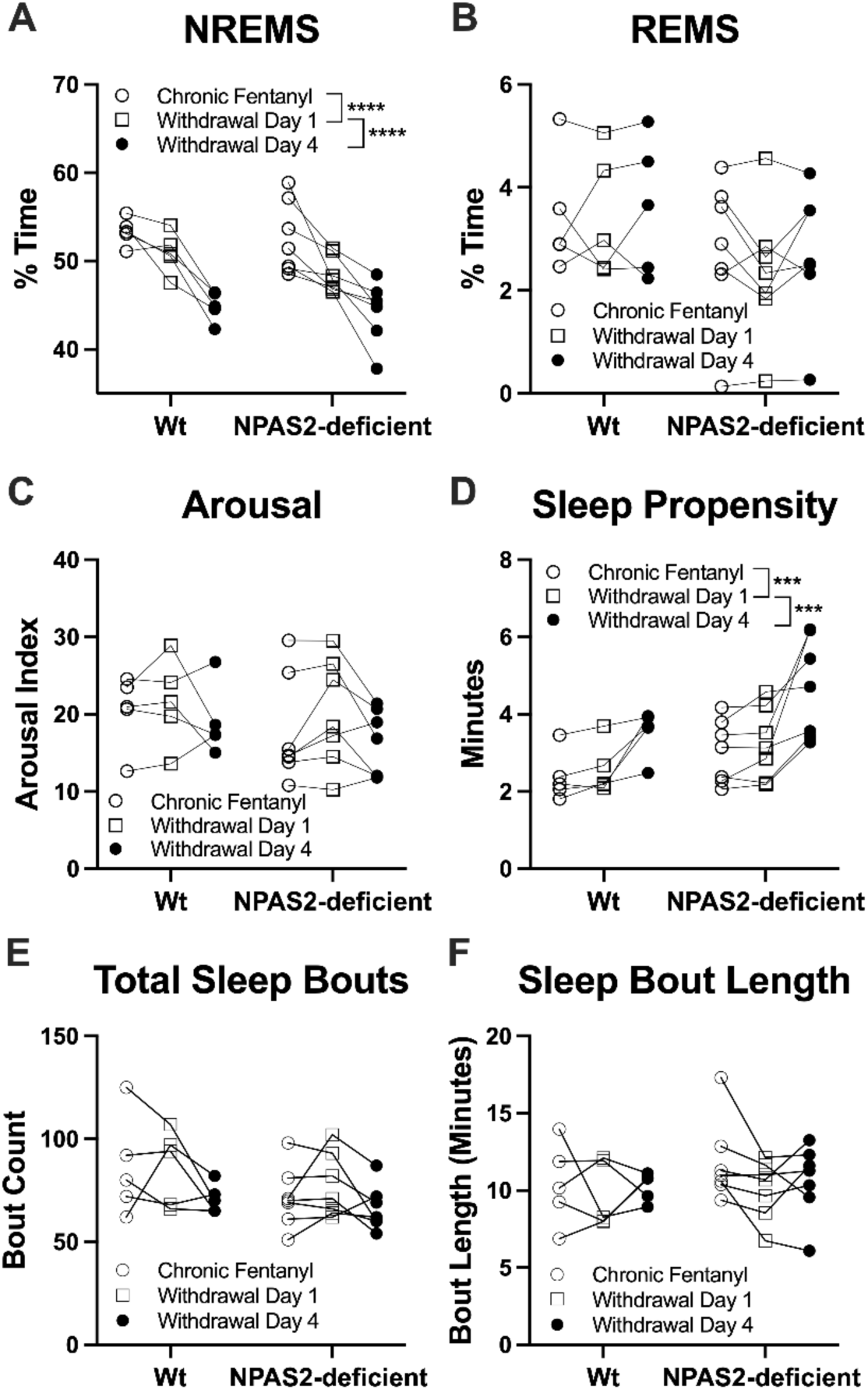
Chronic fentanyl leads to altered sleep and wake states that progressively worsen during fentanyl withdrawal. Sleep-wake states were recorded from wild-type (Wt) and NPAS2- deficient mice after seven days of fentanyl administration and during withdrawal. **A.** Time spent in non-rapid eye movement sleep (NREMS; chronic fentanyl vs. withdrawal day 1, \*\*\*\**p* < 0.0001; withdrawal day 1 vs. withdrawal day 4, \*\*\*\**p* < 0.0001). **B.** Time spent in rapid-eye movement sleep (REMS). **C.** Average number of arousals (transition from sleep to wake state) per hour of sleep. **D.** Time to fall asleep after ≥3 awake epochs (chronic fentanyl vs. withdrawal day 1, \*\*\**p* < 0.001; withdrawal day 1 vs. withdrawal day 4, \*\*\**p* < 0.001). **E.** Total number of sleep bouts over the 24 hrs. **F.** Average length of sleep bouts.

### Diurnal effects of fentanyl administration and withdrawal on sleep and wake in wild-type and NPAS2-deficient mice

To investigate whether sleep-wake cycles were altered by fentanyl, sleep and wake states were analyzed across different times of day. Sleep and wake were largely disrupted during the dark phase of the light-dark cycle in Wt mice, both for acute and chronic fentanyl administration and during withdrawal. Two-way ANOVA indicated a significant effect of time of day on wake (F_(5,84)_ = 22.69; *p* < 0.0001, **Fig. 7A**), NREMS (F_(5,84)_ = 21.95; *p* < 0.0001, **Fig. 7B**), and REMS (F_(5,84)_ = 10.75; *p* < 0.0001, **Fig. 7C**), but no significant effect of fentanyl administration on each of these measures (**Fig. 7A-C**). Similarly, each of these measures varied by time of day (wake: F_(5,84)_ = 57.29; *p* < 0.0001; NREMS: F_(5,84)_ = 50.89; *p* < 0.0001; REMS: F_(5,84)_ = 14.23; *p* < 0.0001) during acute and chronic withdrawal from fentanyl (**Fig. 7A-C**). In addition, significant effects for fentanyl withdrawal were identified for wake (F_(2,84)_ = 3.88; *p* = 0.02; **Fig. 7A**) and NREMS (F_(2,84)_ = 4.48; *p* = 0.01; **Fig. 7B**). Post hoc analyses for fentanyl withdrawal condition revealed a significant increase (*p* < 0.05) in wake on day four of chronic withdrawal compared to acute withdrawal at ZT4 (increased ∼12%; **Fig. 7B**). At ZT20, wake was significantly decreased (*p* < 0.05) on day one of chronic withdrawal (50.38 ± 3.82%) compared to acute withdrawal (62.55 ± 4.41%), suggesting chronic withdrawal alters the diurnal pattern of wakefulness in the early light phase and in the late dark phase (**Fig. 7A**). Fentanyl-induced alterations in REMS were inconsistent by time of day (**Fig. 7C**).

**Fig. 7.**
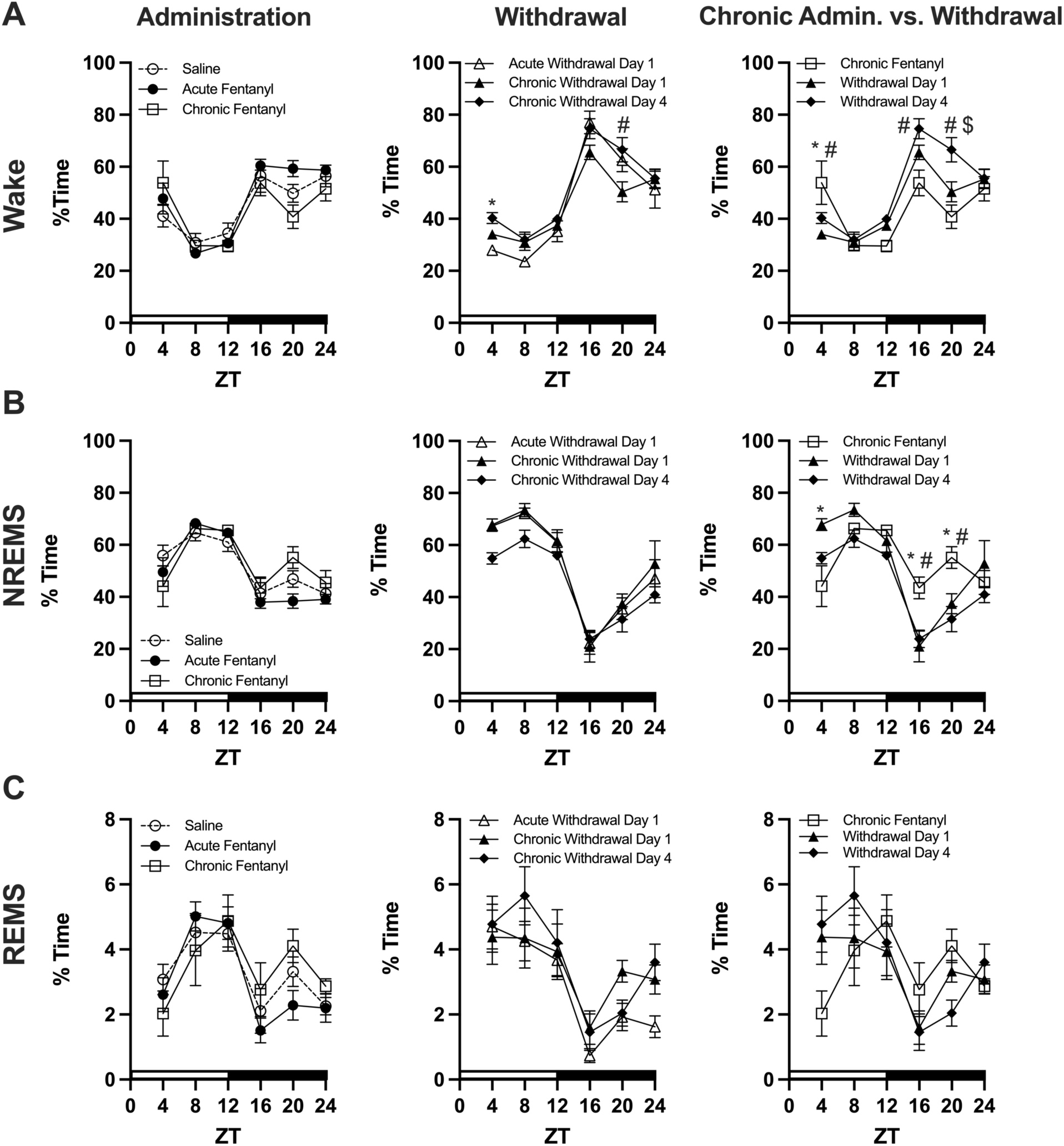
Diurnal effects of fentanyl administration and withdrawal on sleep and wake in wild- type mice. Sleep-wake states across the light-dark cycle in wild-type (Wt) mice. Time spent in wake (**A**), non-rapid eye movement sleep (NREMS, **B**), and rapid eye movement sleep (REMS, **C**) following acute and chronic fentanyl relative to saline (Administration), during fentanyl withdrawal (Withdrawal; # *p* < 0.05, fentanyl withdrawal day one vs. four), and chronic fentanyl administration relative to withdrawal (Chronic Admin. vs. Withdrawal; * *p* < 0.05, chronic fentanyl vs. fentanyl withdrawal day one; # *p* < 0.05, chronic fentanyl vs. withdrawal day four; $ *p* < 0.05, fentanyl withdrawal day one vs. four).

Fentanyl withdrawal led to significant alterations in wake and NREMS that were dependent on time of day. Repeated measures ANOVA identified significant interactions between time of day and fentanyl withdrawal condition for both wake (F_(10,72)_ = 4.279; *p* = 0.0001; **Fig. 7A**) and NREMS (F_(10,72)_ = 4.222; *p* = 0.0001; **Fig. 7B**). Post hoc analyses identified significantly reduced wake on days one and four of withdrawal at ZT4 compared to chronic fentanyl administration (*p* < 0.05). A significant increase in wake was found at ZT16 on day four of withdrawal compared to chronic fentanyl administration (*p* < 0.05), which remained significantly elevated at ZT20 (**Fig. 7A**). Conversely, NREMS was significantly reduced at ZT16 and ZT20 during fentanyl withdrawal on both days one and four (*p* < 0.05; **Fig. 7B**), suggesting the increased wakefulness contributed to reduced NREMS at specific times of day during withdrawal. A main effect for time of day was significant for wake (F_(5,72)_ = 33.37; *p* < 0.0001), NREMS (F_(5,72)_ = 30.16; *p* < 0.0001), and REMS (F_(5,72)_ = 5.37; *p* = 0.0003) during fentanyl withdrawal.

In NPAS2-deficient mice, sleep and wake alterations were also evident following fentanyl administration and withdrawal in a time-of-day dependent manner. Following acute and chronic administration of fentanyl, wake (treatment: F_(2,108)_ = 6.68; *p* = 0.002; and time: F_(5,108)_ = 22.4; *p* < 0.0001) and NREMS (treatment: F_(2,108)_ = 8.28; *p* = 0.0004; and time: F_(5,108)_ = 23.2; *p* < 0.0001) was significantly altered during the light phase of the light-dark cycle. This was evident by an increase in wake following acute fentanyl (saline: 40.16 ± 7.29%; acute fentanyl: 59.80 ± 3.27%; chronic fentanyl: 42.84 ± 2.84%), particularly at ZT4 (*p* < 0.05; **Fig. 8A**), accompanied by a significant decrease (saline: 56.79 ± 6.61%; acute fentanyl: 38.18 ± 3.48%; chronic fentanyl: 54.63 ± 2.41%) in NREMS at ZT4 (*p* < 0.05; **Fig. 8B**). NREMS was also increased by 14% by chronic fentanyl administration (70.63 ± 1.35%) compared to saline (56.70 ± 4.74%) at ZT8 (*p* < 0.05; **Fig. 8B**). No significant effects were observed in REMS between acute and chronic fentanyl administration, only varying by time of day (F_(5,108)_ = 8.68; *p* < 0.0001), with increased REMS during the light, or inactive phase (**Fig. 8C**). Similarly, only main effects for time of day were identified for wake (F_(5,108)_ = 88.37; *p* < 0.0001), NREMS (F_(5,108)_ = 90.43; *p* < 0.0001), and REMS (F_(5,108)_ = 14.05; *p* < 0.0001), during fentanyl withdrawal (**Fig. 8A-C**).

**Fig. 8.**
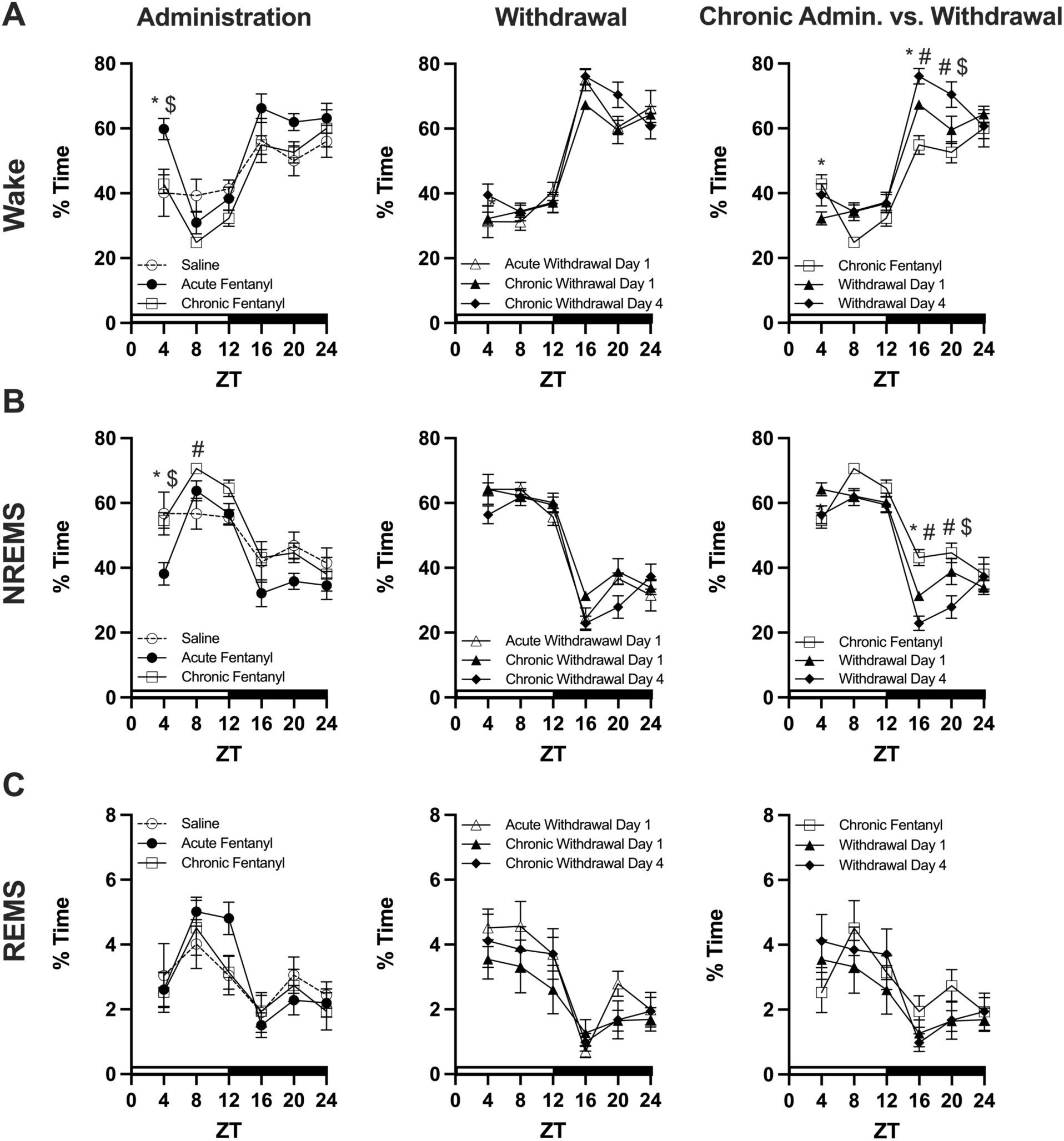
Diurnal effects of fentanyl administration and withdrawal on sleep and wake in NPAS2-deficient mice. Sleep-wake states across the light-dark cycle in NPAS2-deficient mice. Time spent in wake (**A**), non-rapid eye movement sleep (NREMS, **B**), and rapid eye movement sleep (REMS, **C**) following acute and chronic fentanyl relative to saline (Administration; * *p* < 0.05, saline vs. acute fentanyl; $ *p* < 0.05, acute vs. chronic fentanyl; # *p* < 0.05, saline vs. chronic fentanyl), during fentanyl withdrawal (Withdrawal), and chronic fentanyl administration relative to withdrawal (Chronic Admin. vs. Withdrawal; * *p* < 0.05, chronic fentanyl vs. fentanyl withdrawal day one; # *p* < 0.05, chronic fentanyl vs. withdrawal day four; $ *p* < 0.05, fentanyl withdrawal day one vs. four).

Notably, withdrawal from chronic administration of fentanyl led to persistent alterations in sleep and wake in NPAS2-deficient mice. Repeated measures ANOVA revealed significant interactions between fentanyl withdrawal and time of day for wake (F_(10,108)_ = 3.32; *p* = 0.0009; **Fig. 8A**) and NREMS (F_(10,108)_ = 3.33; *p* = 0.0008; **Fig. 8B**). At ZT4, wake was significantly increased following chronic fentanyl (41.10 ± 2.49) relative to withdrawal day one (33.11 ± 2.79) (*p* < 0.05; **Fig. 8A**). Wakefulness was significantly elevated at ZT12 (chronic fentanyl: 29.56 ± 2.20 and withdrawal day four: 39.85 ±1.93) and ZT16 (chronic fentanyl: 53.36 ± 3.76 and withdrawal day four: 76.18 ± 3.18) on day four of withdrawal in NPAS2-deficient mice, accompanied by significant reductions in NREMS at the same times of day (*p* < 0.05; **Fig. 8B**). At ZT4, NREMS was lower for chronic fentanyl administration (ZT4: 56.35 ± 1.99) compared to withdrawal day 1 (ZT4: 63.72 ± 2.90) while at ZT8 and ZT16 NREMS was greater for chronic fentanyl administration (ZT8: 71.80 ± 1.33; ZT16: 44.63 ± 3.22) compared to withdrawal day 1 (ZT8: 62.60 ± 2.23; ZT16: 32.29 ± 1.36; **Fig. 8B**). Similarly, at ZT8, ZT16, and ZT20, NREMS was even lower by withdrawal day 4 (ZT8: 61.31 ± 3.25, ZT16: 22.93 ± 3.01, ZT20: 24.97 ± 3.31) compared to chronic fentanyl administration (ZT8: 71.81 ± 1.33, ZT16: 44.63 ± 3.22, ZT20: 44.68 ± 4.27; **Fig. 8B**). Together, these results demonstrate chronic fentanyl administration led to persistent changes in wake and NREMS through four days of withdrawal in NPAS2-deficient mice.

## Discussion

Intense and severe withdrawal symptoms during abstinence can contribute to the risk of relapse to opioids. Among these symptoms is sleep dysfunction and disrupted circadian rhythms, recognized as a possible hallmark of OUD and opioid dependence (Beswick et al. 2003; Maulik et al. 2002; Lydon-Staley et al. 2017; Eacret et al. 2020). Most patients using prescription opioids or being treated for OUD or dependence with methadone breach the clinical threshold for sleep disorders (Hartwell et al. 2004; Stein et al., 2004). Few studies have characterized the impact of opioids on sleep architecture following acute and chronic opioid administration, as well as the several days into withdrawal. Synthetic opioids have surpassed prescription opioids in rates of use and the cause of drug-related overdose deaths. In the present study, we investigated the impact of the synthetic opioid fentanyl on sleep architecture and the diurnal pattern of sleep-wake states in male mice. In addition, given the recent findings that the circadian transcription factor NPAS2 modulates sleep and drug reward, we assessed whether fentanyl-induced sleep changes were further altered in NPAS2-deficient mice. Overall, our findings demonstrate that both acute and chronic fentanyl lead to changes in sleep architecture, particularly in NREMS, and following chronic administration of fentanyl, alterations in NREMS, accompanied by increased wakefulness and arousal, that persist during withdrawal. Futher, augmented arousal and reduced NREMS was more pronounced in NPAS2-deficient mice, that was specific to transitions between the light and dark phase of the light-dark cycle (*i.e.,* during the early light phase and again during the early dark phase).

Chronic fentanyl administration led to a robust reduction in NREMS that progressively became more severe during days one and four of withdrawal. The progressive loss of NREMS by chronic fentanyl and withdrawal occurred independent of genotype of the mice. Contrary to previous work, fentanyl (citrate) led to increased NREMS and decreased REMS, although these studies only monitored sleep for one hour following an acute fentanyl dose in rats (Montandon and Horner 2019). Here, acute fentanyl administration led to minimal changes in sleep state and wakefulness in wild-type male mice. In contrast, acute fentanyl administration disrupted NREMS and arousal depending on time of day in NPAS2-deficient mice. Acute fentanyl reduced the duration of NREMS in NPAS2-deficient mice and increased the propensity to awaken compared to saline. This reduction in NREMS following fentanyl was more pronounced following either acute or chronic fentanyl administration in NPAS2-deficient mice relative to wild-type controls. Persistent reductions in NREMS were evident on days one and four of withdrawal. Together, our evidence suggests that the circadian transcription factor NPAS2 is involved in mitigating the impact of fentanyl administration on sleep and diurnal regulation of sleep and arousal. Interestingly, chronic fentanyl administration had minimal impacts on sleep and wake immediately following the last injection, suggesting the development of tolerance to opioids and sleep-related changes (Kay 1975).

In addition, NPAS2 may be critical for the regulation of sleep homeostasis in response to perturbations such as stress and sleep deprivation. For example, following sleep deprivation, mice typically experience a rebound of NREMS and REMS (Franken et al. 2006). However, in mice deficient of functional NPAS2, NREMS and REMS rebound following sleep deprivation is significantly blunted (Franken et al. 2006). Moreover, previous work has shown that NPAS2- deficient mice exhibited lower NREMS and REMS in the dark (active) phase of the light-dark cycle (Dudley et al., 2003; Franken et al., 2006). Our findings revealed similar amounts of NREMS and REMS between wild-type and NPAS2-deficient male mice. Nevertheless, our work suggests NPAS2 may be important for mediating the impact of fentanyl on sleep and wake, and possibly, the homeostatic regulation of sleep during withdrawal from opioids. Although the mechanism by which NPAS2 mediates the effects of fentanyl withdrawal on sleep is unknown, the involvement of NPAS2 in the molecular clock is likely (Franken and Dijk 2009) and may be preferentially involved in modulating arousal and wakefulness through actions in the striatum (Becker-Krail et al. 2022; DePoy et al. 2021; Garcia et al. 2000; Luo et al. 2018; Ozburn et al. 2015, 2017; Parekh et al. 2019; Zhang et al. 2021).

NREMS is particularly critical for memory functions, especially declarative memory in humans. Evidence indicates that acute withdrawal impairs working memory in humans (Rapeli et al. 2006), however, studies in rodent models on the effects of acute opioid withdrawal on memory have been inconsistent (Morisot and Contarino 2016; Baidoo et al. 2020). Therefore, altered NREMS during opioid withdrawal may be involved in withdrawal-related impairments in working memory and cognitive function (Rapeli et al. 2006), postulated to contribute to relapse. Disrupted sleep may also contribute to cognitive impairments experienced by people with OUD (Eacret et al. 2020).

While other studies have reported differences in REMS following opioid administration (Eacret et al. 2020; Wang and Teichtahl 2007), our findings suggest that acute and chronic fentanyl administration have minimal effects on REMS in male mice. Additionally, fentanyl has unique pharmacological actions compared to other μ-opioid receptor agonists including morphine, likely contributing to differences between our findings and previous <u>(Kelly et al. 2021)</u>. Additional studies are required to comprehensively understand the relationship between opioids and sleep. Here, we included only male mice, which is a major limitation of our work. Ongoing and future studies include female mice using different models of opioid administration and sleep polysomnography. As OUD continues to rise in females (Barbosa-Leiker et al. 2021), further understanding the sex specific mechanisms underlying the relationships between sleep, circadian rhythms, and opioids is imperative for developing interventions and therapeutics, as opioids differently impact males and females which impact clinical treatment (Huhn et al. 2019). While this study focused on the effects of opioids on sleep, the relationship between sleep and substance use disorders is bidirectional. For instance, chronic sleep deprivation reduced morphine drinking in male and female mice yet developed and maintained morphine conditioned reward (Eacret et al. 2022). Future studies will explicitly investigate sex-specific effects of opioids on sleep and the impact of sleep deprivation on opioid reward. Moreover, considering that fentanyl is often used in combination with other substances (Ciccarone 2019; Peppin et al. 2020), future work should expand into models of polysubstance and sleep.

## Acknowledgments

This study was funded by NHLBI R01HL150432 to R.W.L.

## References

Ahmadi-Soleimani SM, Azizi H, Abbasi-Mazar A (2021) Intermittent REM sleep deprivation attenuates the development of morphine tolerance and dependence in male rats. Neuroscience Letters 748:135735. https://doi.org/10.1016/j.neulet.2021.135735

Baidoo N, Wolter M, Leri F (2020) Opioid withdrawal and memory consolidation. Neuroscience & Biobehavioral Reviews 114:16–24. https://doi.org/10.1016/j.neubiorev.2020.03.029

Barbosa-Leiker C, Campbell ANC, McHugh RK, et al (2021) Opioid Use Disorder in Women and the Implications for Treatment. PRCP 3:3–11. https://doi.org/10.1176/appi.prcp.20190051

Becker-Krail DD, Parekh PK, Ketchesin KD, Yamaguchi S, Yoshin J, Hildebrand MA, Dunham B, Ganapathiraju MK, Logan RW, McClung CA (2022) Circadian transcription factor NPAS2 and NAD+ -dependent deacetylase SIRT1 interact in the mouse nucleus accumbens and regulate reward. Eur J Neurosci 55: 675–693. https://doi.org/10.1111/ejn.15464

Benington JH, Kodali SK, Heller HC (1994) Scoring Transitions to REM Sleep in Rats Based on the EEG Phenomena of Pre-REM Sleep: An Improved Analysis of Sleep Structure. Sleep 17:28– 36. https://doi.org/10.1093/sleep/17.1.28

Beswick T, Best D, Bearn J, Gossop M, Rees S, Strang J (2003) The effectiveness of combined naloxone/lofexidine in opiate detoxification: results from a double-blind randomized and placebo-controlled trial. Am J Addict 12:295–305. https://doi.org/10.1111/j.1521-0391.2003.tb00544.x

Ciccarone D (2019) The Triple Wave Epidemic: Supply and Demand Drivers of the US Opioid Overdose Crisis. Int J Drug Policy 71:183–188. https://doi.org/10.1016/j.drugpo.2019.01.010

Cicero TJ, Ellis MS, Kasper ZA (2020) Polysubstance Use: A Broader Understanding of Substance Use During the Opioid Crisis. Am J Public Health 110:244–250. https://doi.org/10.2105/AJPH.2019.305412

Cicero TJ, Inciardi JA, Muñoz A (2005) Trends in Abuse of OxyContin® and Other Opioid Analgesics in the United States: 2002-2004. The Journal of Pain 6:662–672. https://doi.org/10.1016/j.jpain.2005.05.004

Comer SD, Cahill CM (2019) Fentanyl: Receptor pharmacology, abuse potential, and implications for treatment. Neuroscience & Biobehavioral Reviews 106:49–57. https://doi.org/10.1016/j.neubiorev.2018.12.005

De Andrés I, Caballero A (1989) Chronic morphine administration in cats: Effects on sleep and EEG. Pharmacology Biochemistry and Behavior 32:519–526. https://doi.org/10.1016/0091-3057(89)90191-3

DePoy LM, Becker-Krail DD, Zong W, Petersen K, Shah NM, Brandon JH, Miguelino AM, Tseng GC, Logan RW, McClung CA (2021) J Neurosci 41: 1046-1058. https://doi.org/10.1523/JNEUROSCI.1830-20.2020

Dudley CA, Erbel-Sieler C, Estill SJ, et al (2003) Altered Patterns of Sleep and Behavioral Adaptability in NPAS2-Deficient Mice. Science 301:379–83. https://doi/10.1126/science.1082795

Eacret D, Lemchi C, Caulfield JI, et al (2022) Chronic Sleep Deprivation Blocks Voluntary Morphine Consumption but Not Conditioned Place Preference in Mice. Front Neurosci 16:836693. https://doi.org/10.3389/fnins.2022.836693

Eacret D, Veasey SC, Blendy JA (2020) Bidirectional Relationship between Opioids and Disrupted Sleep: Putative Mechanisms. Mol Pharmacol 98:445–453. https://doi.org/10.1124/mol.119.119107

Fathi HR, Yoonessi A, Khatibi A, et al (2020) Crosstalk between Sleep Disturbance and Opioid Use Disorder: A Narrative Review. Addict Health 12:140–158. https://doi.org/10.22122/ahj.v12i2.249

Franken P, Dijk D-J (2009) Circadian clock genes and sleep homeostasis. European Journal of Neuroscience 29:1820–1829. https://doi.org/10.1111/j.1460-9568.2009.06723.x

Franken P, Dudley CA, Estill SJ, et al (2006) NPAS2 as a transcriptional regulator of non-rapid eye movement sleep: Genotype and sex interactions. Proc Natl Acad Sci U S A 103:7118–7123. https://doi.org/10.1073/pnas.0602006103

Garcia JA, Zhang D, Estill SJ, et al (2000) Impaired Cued and Contextual Memory in NPAS2- Deficient Mice. Science 288:2226–2230. https://doi.org/10.1126/science.288.5474.2226

Gladden RM (2016) Fentanyl Law Enforcement Submissions and Increases in Synthetic Opioid– Involved Overdose Deaths — 27 States, 2013–2014. MMWR Morb Mortal Wkly Rep 65:. https://doi.org/10.15585/mmwr.mm6533a2

Hartwell EE, Pfeifer JG, McCauley JL, Moran-Santa Maria M, Back SE (2014) Sleep disturbances and pain among individuals with prescription opioid dependence. Addict Behav 39: 1537–1542. https://doi:10.1016/j.addbeh.2014.05.025

Huhn AS, Berry MS, Dunn KE (2019) Review: Sex-based Differences in Treatment Outcomes for Persons with Opioid Use Disorder. Am J Addict 28:246–261. https://doi.org/10.1111/ajad.12921

Kay DC (1975) Human sleep during chronic morphine intoxication. Psychopharmacologia 44:117–124. https://doi.org/10.1007/BF00420997

Kay DC, Eisenstein RB, Jasinski DR (1969) Morphine effects on human REM state, waking state and NREM sleep. Psychopharmacologia 14:404–416. https://doi.org/10.1007/BF00403581

Kelly E, Sutcliffe K, Cavallo D, et al (2021) The anomalous pharmacology of fentanyl. British Journal of Pharmacology 1:. https://doi.org/10.1111/bph.15573

Khazan N, Colasanti B (1972) Protracted rebound in rapid movement sleep time and electroencephalogram voltage output in morphine-dependent rats upon withdrawal. J Pharmacol Exp Ther 183:23–30

Lewis SA, Oswald I, Evans JI, et al (1970) Heroin and human sleep. Electroencephalography and Clinical Neurophysiology 28:374–381. https://doi.org/10.1016/0013-4694(70)90230-0

Logan RW, Williams WP, McClung CA (2014) Circadian rhythms and addiction: Mechanistic insights and future directions. Behav Neurosci 128:387–412. https://doi.org/10.1037/a0036268

Luo YJ, Li YD, Wang L, Yang SR, Yuan XS, Wang J, Cherasse Y, Lazarus M, Chen JF, Qu WM, Huang ZL (2018) Nucleus accumbens controls wakefulness by a subpopulation of neurons expressing dopamine D1 receptors. Nat Commun 9: 1576. https://doi:10.1038/s41467-018-03889-3

Lydon-Staley DM, Cleveland HH, Huhn AS, et al (2017) Daily sleep quality affects drug craving, partially through indirect associations with positive affect, in patients in treatment for nonmedical use of prescription drugs. Addict Behav 65:275–282. https://doi.org/10.1016/j.addbeh.2016.08.026

Maulik PK, Tripathi BM, Pal HR (2002) Coping behaviors and relapse participants in opioid dependence: a study from North India. J Subst Abuse Treat. 22: 135–140. https://doi:10.1016/s0740-5472(02)00225-8

McCullough KM, Missig G, Robble MA, Foib AR, Wells AM, Hartmann J, Anderson KJ, Neve RL, Nestler EJ, Ressler KJ, Carlezon Jr WA (2021) Nucleus accumbens medium spiny neuron subtypes differentially regulate stress-associated alterations in sleep architecture. Biol Psychiatry 89: 1138–1149. https://doi:10.1016/j.biopsych.2020.12.030

Montandon G, Horner RL (2019) Electrocortical changes associating sedation and respiratory depression by the opioid analgesic fentanyl. Sci Rep 9:14122. https://doi.org/10.1038/s41598-019-50613-2

Morisot N, Contarino A (2016) The CRF1 and the CRF2 receptor mediate recognition memory deficits and vulnerability induced by opiate withdrawal. Neuropharmacology 105:500–507. https://doi.org/10.1016/j.neuropharm.2016.02.021

O’Donnell BJ, Guo L, Ghosh S, et al (2019) Sleep phenotype in the Townes mouse model of sickle cell disease. Sleep Breath 23:333–339. https://doi.org/10.1007/s11325-018-1711-x

O’Donnell JK (2017) Deaths Involving Fentanyl, Fentanyl Analogs, and U-47700 — 10 States, July–December 2016. MMWR Morb Mortal Wkly Rep 66:.https://doi.org/10.15585/mmwr.mm6643e1

Orr WC, Stahl ML (1978) Sleep Patterns in Human Methadone Addiction. Addiction 73:311–315. https://doi.org/10.1111/j.1360-0443.1978.tb00158.x

Oswald I (1969) Sleep, dreaming and drugs. Proc R Soc Med 62:151–153

Ozburn AR, Falcon E, Twaddle A, et al (2015) Direct regulation of diurnal Drd3 expression and cocaine reward by NPAS2. Biol Psychiatry 77:425–433. https://doi.org/10.1016/j.biopsych.2014.07.030

Ozburn AR, Kern J, Parekh PK, Logan RW, Liu Z, Falcon E, Becker-Krail D, Purohit K, Edgar NM, Huang Y, McClung CA (2017) NPAS2 regulation of anxiety-like behavior and GABAA receptors. Front Mol Neurosci. 10:360. https://doi:10.3389/fnmol.2017.00360

Parekh PK, Logan RW, Ketchesin KD, Becker-Krail D, Shelton MA, Hildebrand MA, Barko K, Huang YH, McClung CA. (2019) Cell-type-specific regulation of nucleus accumbens synaptic plasticity and cocaine reward sensitivity by the circadian protein, NPAS2. J Neurosci. 39: 4657-4667. https://doi.org/10.1523/JNEUROSCI.2233-18.2019

Peppin JF, Raffa RB, Schatman ME (2020) The Polysubstance Overdose-Death Crisis. J Pain Res 13:3405–3408. https://doi.org/10.2147/JPR.S295715

Rapeli P, Kivisaari R, Autti T, et al (2006) Cognitive function during early abstinence from opioid dependence: a comparison to age, gender, and verbal intelligence matched controls. BMC Psychiatry 6:9. https://doi.org/10.1186/1471-244X-6-9

Shaw IR, Lavigne G, Mayer P, Choinière M (2005) Acute Intravenous Administration of Morphine Perturbs Sleep Architecture in Healthy Pain-Free Young Adults: a Preliminary Study. Sleep 28:677–682. https://doi.org/10.1093/sleep/28.6.677

Smith MT, Mun CJ, Remeniuk B, et al (2020) Experimental sleep disruption attenuates morphine analgesia: findings from a randomized trial and implications for the opioid abuse epidemic. Sci Rep 10:20121. https://doi.org/10.1038/s41598-020-76934-1

Stein MD, Herman DS, Bishop S, Lassor JA, Weinstock M, Anthony J, Andersen BJ (2004) Sleep disturbances among methadone maintained patients. J Subst Abuse Treat 26: 175–180. https://doi:10.1016/S0740-5472(03)00191-0

Tagaito Y, Polotsky VY, Campen MJ, et al (2001) A model of sleep-disordered breathing in the C57BL/6J mouse. Journal of Applied Physiology 91:2758–2766. https://doi.org/10.1152/jappl.2001.91.6.2758

Tripathi R, Rao R, Dhawan A, et al (2020) Opioids and sleep – a review of literature. Sleep Medicine 67:269–275. https://doi.org/10.1016/j.sleep.2019.06.012

Veasey SC, Yeou-Jey H, Thayer P, Fenik P (2004) Murine Multiple Sleep Latency Test: Phenotyping Sleep Propensity in Mice. Sleep 27:388–393. https://doi.org/10.1093/sleep/27.3.388

Wang D, Teichtahl H (2007) Opioids, sleep architecture and sleep-disordered breathing. Sleep Medicine Reviews 11:35–46. https://doi.org/10.1016/j.smrv.2006.03.006

Zhang R, Manza P, Tomasi D, Kim SW, Shokri-Kojori E, Demiral SB, Kroll DS, Feldman DE, McPherson KL, Biesecker CL, Wang GJ, Volkow ND (2021) Dopamine D1 and D2 receptors are distinctly associated with rest-activity rhythms and drug reward. J Clin Invest 131: e149722. https://doi:10.1172/JCI149722

